# Multiplexed functional assessments of *MYH7* variants in human cardiomyocytes at scale

**DOI:** 10.1101/2023.07.28.551073

**Authors:** Clayton E. Friedman, Shawn Fayer, Sriram Pendyala, Wei-Ming Chien, Linda Tran, Leslie S. Chao, Ashley McKinstry, Dania Ahmed, Alexander Loiben, Stephen D. Farris, April Stempien-Otero, Erica Jonlin, Charles E. Murry, Lea M. Starita, Douglas M. Fowler, Kai-Chun Yang

**Affiliations:** Institute for Stem Cell and Regenerative Medicine, University of Washington, School of Medicine, Seattle, WA 98109, USA; Center for Cardiovascular Biology, University of Washington, Seattle, WA 98109, USA; Department of Medicine/Cardiology, University of Washington, Seattle, WA 98109, USA; Department of Genome Sciences, University of Washington, Seattle, WA 98195, USA; Cardiology/Hospital Specialty Medicine, VA Puget Sound HCS, Seattle, WA 98108, USA; Department of Pathology, University of Washington, Seattle, WA 98109, USA; Department of Bioengineering, University of Washington, Seattle, WA 98195, USA; Brotman Baty Institute for Precision Medicine, Seattle, WA 98195, USA

**Keywords:** *MYH7*, cardiomyocytes, CRaTER, hypertrophic cardiomyopathy, hypertrophy, survival, stem cells, hiPSCs, VAMP-seq, DMS

## Abstract

**Background:** Single, autosomal-dominant missense mutations in *MYH7*, which encodes a sarcomeric protein (MHC-β) in cardiac and skeletal myocytes, are a leading cause of hypertrophic cardiomyopathy and are clinically-actionable. However, ∼75% of *MYH7* variants are of unknown significance (VUS), causing diagnostic challenges for clinicians and emotional distress for patients. Deep mutational scans (DMS) can determine variant effect at scale, but have only been utilized in easily-editable cell lines. While human induced pluripotent stem cells (hiPSCs) can be differentiated to numerous cell types that enable the interrogation of variant effect in a disease-relevant context, DMS have not been executed using diploid hiPSC derivates. However, CRaTER enrichment has recently enabled the pooled generation of a saturated five position *MYH7* variant hiPSC library suitable for DMS for the first time.

**Results:** As a proof-of-concept, we differentiated this *MYH7* variant hiPSC library to cardiomyocytes (hiPSC-CMs) for multiplexed assessment of MHC-β variant abundance by massively parallel sequencing (VAMP-seq) and hiPSC-CM survival. We confirm MHC-β protein loss occurs in a failing human heart with a pathogenic *MYH7* mutation. We find the multiplexed assessment of MHC-β abundance and hiPSC-CM survival both accurately segregate all pathogenic variants from synonymous controls. Overall, functional scores of 68 amino acid substitutions across these independent assays are ∼50% consistent.

**Conclusions:** This study leverages hiPSC differentiation into disease-relevant cardiomyocytes to enable multiplexed assessments of *MYH7* missense variants at scale for the first time. This proof-of-concept demonstrates the ability to DMS previously restricted, clinically-actionable genes to reduce the burden of VUS on patients and clinicians.

## Background

While every possible single nucleotide variant (SNV) compatible with life is likely represented within the ∼8 billion genomes of *Homo sapiens* [1], the majority of variant effects are unknown [2]. Accurate variant effect data provides functional evidence that is integrated with other clinical evidence to classify variants of unknown significance (VUS) following standard guidelines [3]. VUS cause diagnostic challenges for clinicians and emotional distress for patients. Low allele frequency in the general population can suggest variant pathogenicity, however, this evidence is limited by sparse genome sampling of rare variants. Computational approaches have been developed to overcome rare variation in order to predict variant effect across the human exome, however, inconsistent predictive accuracy between and within genes can limit their general utility [4]. As a result, functional data is particularly useful for variant classification in clinically-actionable genes identified by the American College of Medical Genetics/Genomics [5]. VUS classification in such clinically-actionable genes directly guides clinical management to improve patient care.

Deep mutational scanning (DMS) simultaneously measures the effect of hundreds to thousands of genetic variants to provide functional evidence for variant classification at scale [6, 7]. However, DMS have not been executed in many clinically-actionable genes because their expression and disease manifestation occur in cell types that are difficult to mutagenize and/or phenotype at scale. The majority of clinically-actionable genes encode cardiac or neuronal proteins [5] that would likely require the native intracellular context, including diverse interacting proteins and regulatory chaperones, in order to accurately determine variant effect. Human induced pluripotent stem cells (hiPSCs) have the capacity to differentiate to nearly any cell type in the body and numerous protocols enable directed differentiation of hiPSCs to specific, disease-relevant cell types [8, 9]. For example, hiPSC-CMs are a powerful tool to assess variant effect in clinically-actionable genes and have provided functional evidence for VUS classification [10-16], however, low-throughput variant generation and assessment methods are outpaced by the discovery of novel VUS [17]. To date, DMS has not been performed in diploid hiPSC derivatives due to low genome-editing rates that have prevented the generation of large variant libraries [18].

To enable DMS in hiPSC derivatives, we leveraged a method we developed called <u>CR</u>ISPR<u>a</u> On-<u>T</u>arget <u>E</u>diting <u>R</u>etrieval (CRaTER) that allowed the pooled generation of a variant library in hiPSCs for the first time [19]. This library of isogenic gene-edited hiPSCs targeted the endogenous *MYH7* locus with a single, heterozygous missense variant per cell to mutagenize five amino acid positions of *MYH7* to saturation [19]. *MYH7* encodes a sarcomeric thick filament protein (MHC-β) expressed in cardiac and skeletal myocytes that interacts with actin to generate contractile force. Single, autosomal-dominant missense mutations in *MYH7* are a leading cause of hypertrophic cardiomyopathy (HCM) [20, 21]. All pathogenic/likely pathogenic (P/LP) *MYH7* variants identified from genetic testing are clinically actionable [22] because early HCM diagnosis and intervention can help lower the risk of sudden cardiac death [22, 23].

Here we describe, to our knowledge, the first DMS proof-of-concept in a diploid hiPSC derivative. We differentiated *MYH7* missense variant library hiPSCs to cardiomyocytes in bulk and performed multiple, independent multiplexed assays to determine variant effects. Pathogenic variants in many genes can disrupt protein stability and reduce abundance [24-26], thus, we optimized a FACS-based assay to simultaneously measure MHC-β abundance from the pooled library of *MYH7* missense variant hiPSC-CMs. In addition, our lab [27] and others [11] have shown that pathogenic *MYH7* variant hiPSC-CMs have reduced survival relative to synonymous controls. We find that both the multiplexed assessment of MHC-β abundance and survival in the *MYH7* variant library hiPSC-CMs correctly segregated all tested pathogenic/likely pathogenic (3/3) and synonymous (3/3) variants, providing functional variant effect data for 68 amino acid substitutions. We also find that variants with MHC-β loss also have significantly increased cell size, a hallmark of HCM [27, 28]. In addition, 3/4 *MYH7* VUS examined were functionally abnormal across both assays, suggesting they are likely deleterious.

Altogether, this study leverages the differentiation capacity of hiPSC variant libraries generated by CRaTER in combination with multiplexed assays to determine variant effect at scale, and, in doing so, establishes a framework for the expansion of DMS into clinically-actionable genes and disease-relevant cell types.

## Results

### Human cardiomyocytes with an HCM-causing *MYH7* missense mutation express less MHC-β protein

Pathogenic missense variants in numerous genes are known to disrupt protein stability and as a result, reduce protein abundance [24, 25]. We hypothesized that pathogenic *MYH7* missense variants would similarly cause decreased MHC-β abundance and that this phenotype could be leveraged to functionally assess the effect of *MYH7* missense variants.

To test this hypothesis, we first examined MHC-β protein expression in human heart samples and in human induced pluripotent stem cell-derived cardiomyocytes (hiPSC-CMs) (**Fig. 1**). Human myocardium samples were collected from unrelated individuals without heart failure (non-failing; MYH7^+/+^) or with heart failure (failing; MYH7^E848G/+^) and protein lysates were probed for expression of MHC-β and cardiac troponin I (cTnI) by Western blot (**Fig. 1A-D**; **Supp. Fig. S1**). The patient with the failing heart has a heterozygous pathogenic/likely pathogenic (P/LP) *MYH7* missense variant (c.2543A>G; p.Glu848Gly) known to cause hypertrophic cardiomyopathy (HCM) [10]. MHC-β expression was markedly reduced (average 71% lower) in failing versus non-failing myocardium (**Fig. 1C**), despite similar cTnI expression between non-failing and failing conditions suggesting that the sampled myocardium was composed of similar proportions of cardiomyocytes relative to non-myocyte cell types (**Fig. 1D**). These data reveal a novel link between MHC-β depletion and a pathogenic *MYH7* mutation in a patient heart.

**Fig. 1.**
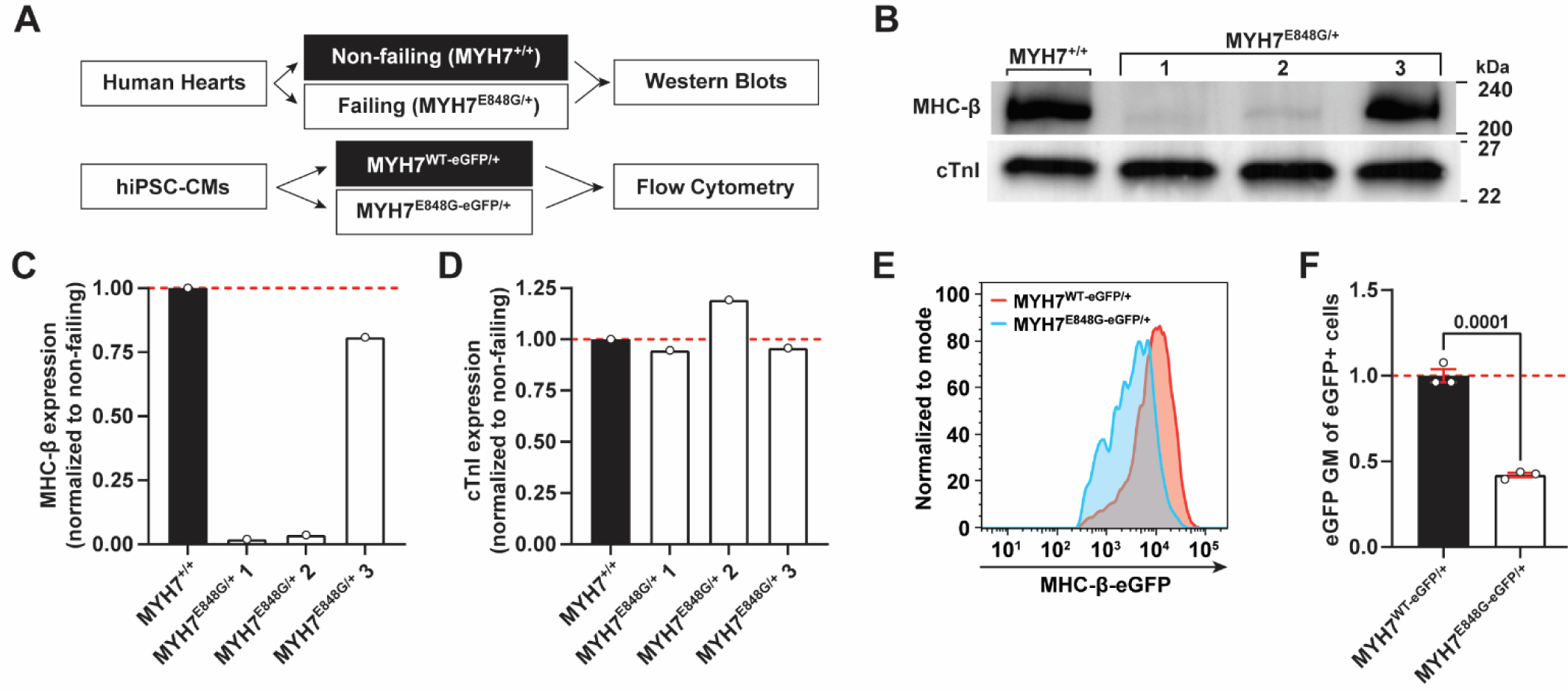
Sarcomeric protein abundance in human hearts and in isogenic hiPSC-CMs. **A** Experimental strategies to quantify β myosin heavy chain (MHC-β) protein abundance in human hearts and isogenic human induced pluripotent stem cell-derived cardiomyocytes (hiPSC-CMs). **B** Representative Western blots for MHC-β (220 kDa) and cardiac troponin I (cTnI; 24 kDa) protein expression in non-failing (MYH7^+/+^, *n* = 1 heart sample) and failing (MYH7^E848G/+^, *n* = 3 samples from the same patient) human heart samples. **C**-**D** Quantifications of MHC-β and cTnI protein expression from **B** normalized to non-failing. **E** Representative flow cytometry histogram overlaying eGFP intensities of age-matched MYH7^WT-eGFP/+^ (red) and MYH7^E848G-eGFP/+^ (blue) isogenic cardiomyocytes. WT = wildtype. **F** Quantifications of eGFP geometric means (GM) from **E** normalized to MYH7^WT-eGFP/+^ (*n* = 3 independent replicates). Error bars indicate standard error of the mean. *P* values calculated using unpaired *t* test; *p* < 0.05 considered significant. Dashed red line in **C**, **D**, and **F** indicates normalized mean. See **Supplemental Figure S1** for complete and stain-free Western blots.

To rule out the contribution of patient-specific disease modifiers to the MHC-β depletion phenotype, we examined isogenic, gene-edited hiPSC cell lines with a heterozygous, C-terminus eGFP-tagged *MYH7* allele to enable MHC-β-eGFP intensity quantification by flow cytometry as a proxy for MHC-β abundance [27]. MHC-β-eGFP quantification in these hiPSC-CMs revealed a significant reduction in eGFP intensity for MYH7^E848G-eGFP/+^ hiPSC-CMs (42% lower) relative to isogenic MYH7^WT-eGFP/+^ hiPSC-CMs lacking the pathogenic mutation (**Fig. 1E-F**). Taken together, these data demonstrate that a missense mutation in *MYH7* can cause MHC-β protein depletion, both in the human heart *in vivo* and in isogenic hiPSC-CMs *in vitro*.

### A multiplexed assessment of MHC-β abundance in hiPSC-CMs containing a library of heterozygous *MYH7* missense variants accurately predicts pathogenicity

We hypothesized that MHC-β protein depletion could be a common phenotype among pathogenic missense variants. To test this hypothesis, we used a previously generated, biallelically-edited hiPSC library containing 113 heterozygous *MYH7* missense variants encoding 78 different amino acid substitutions [19]. Within each of these diploid cells (generated on a healthy WTC11 background) there is one WT *MYH7* allele C-terminally tagged with eGFP while the other *MYH7* allele contains a single *MYH7* variant similarly tagged with mTagBFP2. Tagging both *MYH7* alleles with different fluorescent proteins was originally designed to enable enrichment of heterozygously gene-edited cells using CRaTER [19]. In this study, we leveraged these MHC-β fluorescent protein fusions to enable simultaneous assessment of MHC-β-based variant abundance by massively parallel sequencing (VAMP-seq [26]) in bulk differentiated hiPSC-CMs to examine MHC-β abundance in a multiplexed fashion.

To this end, we first examined eGFP and mTagBFP2 intensity in pooled *MYH7* variant library hiPSC-CMs across a time course of cardiomyocyte maturation (25, 28, 32, and 47 days after the onset of differentiation) using flow cytometry to identify the optimum timepoint to perform VAMP-seq (**Fig. 2A**). We observed the emergence of a mTagBFP2^low^/eGFP^+^ population in *MYH7* variant library hiPSC-CMs that was not present in age-matched MYH7^WT-eGFP/WT-mTagBFP2^ (WT) hiPSC-CMs. Given our finding that cardiomyocytes with the p.Glu848Gly (P/LP) *MYH7* mutation have MHC-β protein loss (**Fig. 1**), we reasoned that there would be an enrichment for pathogenic *MYH7* variants in the mTagBFP2^low^/eGFP^+^ population relative to the mTagBFP2^high^/eGFP^+^ population. We tested this theory directly by harvesting T47 *MYH7* variant library hiPSC-CMs, performing fluorescence-activated cell sorting (FACS) of eGFP^+^ cardiomyocytes into bins on the basis of variable mTagBFP2 intensity (low, mid, and high), preparing *MYH7* amplicon libraries for next generation sequencing (NGS) of each bin, and calculating variant abundance scores for each *MYH7* missense variant in the library (**Fig. 2B**). mTagBFP2**^-^** cells were excluded from sorting to be certain only biallelically-edited cells were analyzed. This experiment was repeated three times using independent cardiac differentiations, however, due to insufficient cell numbers, corresponding bins from the second and third replicates were combined for NGS after sorting. Preliminary comparative analysis of mTagBFP2 low, mid, and high sorting gates revealed a significant decrease in mTagBFP2 intensity in the low bin relative to the mid and high bins, as expected (**Supp. Fig. S2A-B**). In addition, we observed a slight, yet significant increase in FSCA, a proxy for cell size, which is a hallmark phenotype of HCM [27-29], in the low mTagBFP2 bin relative to the mid and high bins (**Supp. Fig. S2C-D**). This novel correlation between decreased MHC-β protein and increased cell size (**Supp. Fig. S2E**) supports our hypothesis that mTagBFP2^low^/eGFP^+^ hiPSC-CMs are more likely to be pathogenic.

**Fig. 2.**
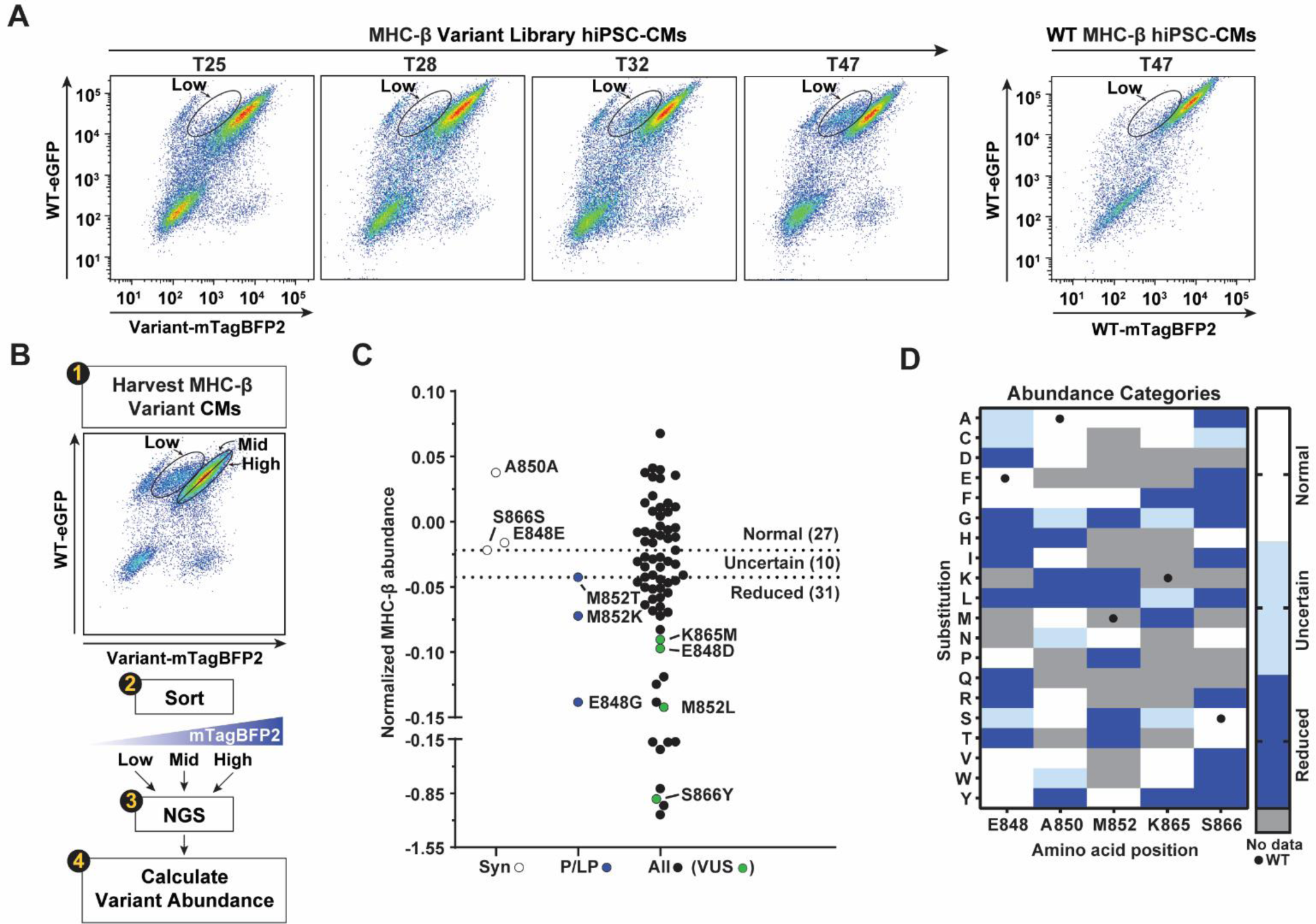
MHC-β abundance in a library of heterozygous *MYH7* missense variant hiPSC-CMs. **A** Live flow cytometry time course analysis of Variant-MHC-β-mTagBFP2 (x-axis) and WT-MHC-β-eGFP (y-axis) expression from a single, cardiac-directed differentiation of biallelically-edited hiPSCs containing a library of heterozygous *MYH7* variants (MYH7^WT-eGFP/Variant-mTagBFP2^; generated in Friedman *et al.* [19]), left, or age-matched, wildtype (WT) cardiomyocytes (MYH7^WT-eGFP/WT-mTagBFP2^), right. T = days after the activation of differentiation. Low = mTagBFP2^low^/eGFP^+^ cells. **B** Experimental strategy to quantify MHC-β-mTagBFP2 abundance in MYH7^WT-eGFP/Variant-mTagBFP2^ hiPSC-CMs using variant abundance by massively parallel sequencing (VAMP-seq [26]). Low, Mid, and High = varying degrees of mTagBFP2 intensity. NGS = next generation sequencing. **C** Mean normalized MHC-β-mTagBFP2 abundance scores for synonymous (Syn; white), pathogenic/likely pathogenic (P/LP; blue), and all variants (black) (*n* = 2 independent replicates). Variants of unknown significance (VUS) scores in ‘All’ column = green dots. Upper dotted line = lowest ‘Normal’ abundance score; lower dotted line = highest ‘Reduced’ abundance score; scores between ‘Normal’ and ‘Reduced’ are ‘Uncertain’. Note broken y-axis. **D** Position map displaying categorical MHC-β-mTagBFP2 abundance scores for each mutagenized residue (vertical) relative to amino acid substitutions (horizontal). Grey = no data. Black dots = WT residues. See **Supplemental Figure S2** for pooled analyses of *MYH7* variant hiPSC-CM library mTagBFP2 intensity and FSCA. See **Supplemental Figure S3** for VAMP-seq replicate data.

In total, these experiments generated functional scores for 68 amino acid substitutions that were normalized to three synonymous (Syn) variants (p.Glu848=, p.Ala850=, and p.Ser866=) assumed to not impact protein stability and/or abundance because they encode wildtype residues and are unlikely to affect splicing given they are at >110 nucleotides form the nearest intron/exon boundary. Normalized MHC-β abundance scores between independent replicates were highly consistent (R^2^ = 0.9447), indicating MHC-β-based VAMP-seq detects a reproducible phenotype (**Supp. Fig. S3**). 27/68 (39.7%) amino acid substitutions resulted in abundances greater than the lowest Syn variant and were considered functionally Normal (**Fig. 2C-D**). All tested pathogenic/likely pathogenic (P/LP) substitutions (p.Glu848Gly [P/LP], p.Met850Thr [P/LP], and p.Met852Lys [LP]) resulted in reduced MHC-β abundance relative to Syn variants (**Fig. 2C-D**). 31/68 (45.6%) amino acid substitutions resulted in abundances lower than the highest P/LP variant and were considered functionally Reduced (**Fig. 2C-D**). Many amino acid substitutions at the p.Ser866 residue resulted in severely reduced MHC-β abundances (**Supp. Fig. S3B-D**). 10/68 (14.7%) amino acid substitutions resulted in abundances lower than the lowest Syn variant yet higher than the highest P/LP variant and were considered functionally Uncertain (**Fig. 2C-D**). Together, multiplexed, MHC-β-based VAMP-seq in hiPSC-CMs accurately segregates all tested P/LP variants (3/3) and VUS (4/4) from synonymous variants (3/3), suggesting this assay could be a useful approach for classifying newly observed *MYH7* variants in the future.

### Examination of MHC-β abundance in clonal heterozygous *MYH7* variant hiPSC-CMs recapitulates multiplexed VAMP-seq

In order to validate the multiplexed VAMP-seq findings, clonal, biallelically-edited *MYH7* variant library hiPSCs previously isolated by our group [19] were individually differentiated to cardiomyocytes for MHC-β abundance phenotyping (**Fig. 3**; **Supp. Fig. S4**). Relative to T47 MYH7^WT-eGFP/WT-mTagBFP2^ hiPSC-CMs (WT), MYH7^WT-eGFP/S866F-mTagBFP2^ and MYH7^WT-eGFP/S866W -mTagBFP2^ hiPSC-CMs showed significantly reduced mTagBFP2 expression (average 68% and 87% lower, respectively), whereas MYH7^WT-eGFP/E848E-mTagBFP2^ hiPSC-CMs showed an insignificant 8% reduction, as expected for a synonymous variant (**Fig. 3A-B**). MYH7^WT-eGFP/M852T-mTagBFP2^ (P/LP), ^MYH7WT-eGFP/K865M-mTagBFP2^ (VUS), and MYH7^WT-eGFP/E848D-mTagBFP2^ (VUS) hiPSC-CMs showed a robust, albeit insignificant, reduction in mTagBFP2 expression relative to WT (average 45%, 24%, and 33% lower, respectively) (**Fig. 3B**). Earlier examination of mTagBFP2 intensity in the same clonal hiPSC-CMs at T30 reveals similar trends with a more modest loss of mTagBFP2 relative to the T47 timepoint, reflecting the progressive nature of mTagBFP2 depletion (**Fig. 2A**). Importantly, mTagBFP2 intensities measured from clonal cell lines are highly correlated with corresponding, age-matched VAMP-seq abundance scores (R^2^ = 0.8233), validating that pooled differentiation and multiplexed assessment of variant effect yield comparable results to individual assessment (**Fig. 3C**). Further examination of eGFP intensity in the same clonal hiPSC-CMs shows a consistent, albeit insignificant, 30-40% decrease across all genotypes (save MYH7^WT-eGFP/E848E-mTagBFP2^) relative to WT, suggesting a potential effect of Variant-MHC-β-mTagBFP2 depletion is the partial loss of WT-MHC-β-eGFP (**Fig. 3D**).

**Fig. 3.**
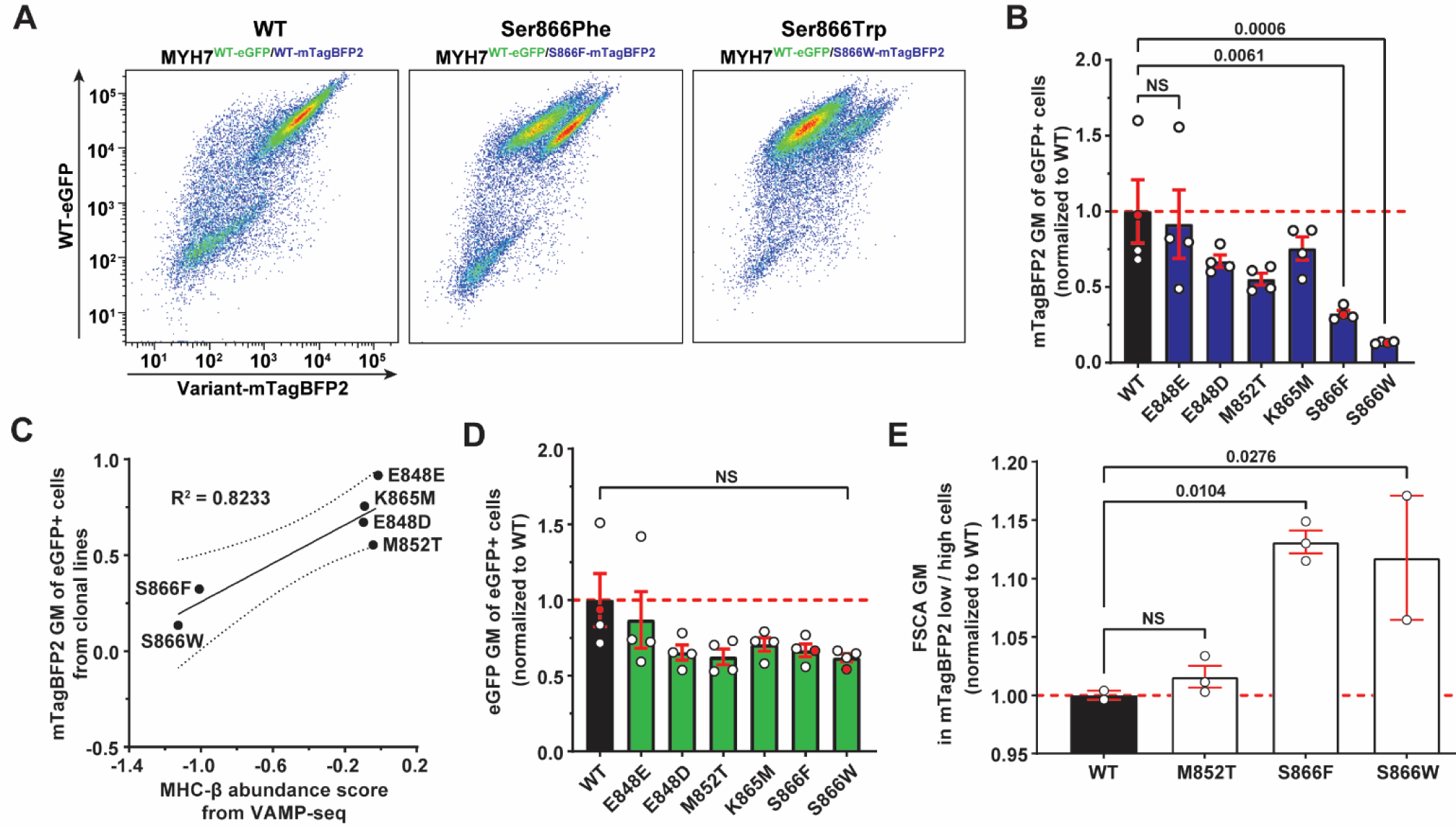
MHC-β abundance in clonal heterozygous *MYH7* variant hiPSC-CMs. **A** Representative flow cytometry plots of Variant-MHC-β-mTagBFP2 (x-axis) and WT-MHC-β-eGFP (y-axis) expression in biallelically-edited T47 hiPSC-CMs containing a single heterozygous *MYH7* variant (left to right; MYH7^WT-eGFP/WT-mTagBFP2^ (WT), MYH7^WT-eGFP/S866F-mTagBFP2^, and MYH7^WT-eGFP/S866W-mTagBFP2^). **B** Quantifications of mTagBFP2 geometric means (GM) from **A** normalized to WT (*n* = 4 independent replicates). **C** Simple linear regression comparing mean normalized MHC-β abundance scores from VAMP-seq (x-axis; Fig. 2I) to mean normalized mTagBFP2 GMs from corresponding clonal heterozygous *MYH7* variant hiPSC-CMs (y-axis). Solid line = best-fit line. Dotted lines = 95% confidence intervals. **D** Quantifications of eGFP GMs from **A** normalized to WT (*n* = 4 independent replicates). **E** Quantifications of FSCA GM in mTagBFP2 low versus high hiPSC-CMs plated at high density, normalized to the corresponding measure in MYH7^WT-eGFP/WT-mTagBFP2^ hiPSC-CMs (*n* = 2-3 independent replicates). Dashed red lines in **B** and **D** indicate normalized WT means and red dots correspond to plots shown in **A**. Error bars indicate standard error of the mean. *P* values calculated using repeated measure one-way ANOVA (**B** and **D**) or one-way ANOVA (**E**); *p* < 0.05 considered significant; not significant (NS). See **Supplemental Figure S4** for confocal microscopy of clonal heterozygous *MYH7* variant hiPSC-CMs.

These results clearly demonstrate a previously unknown correlation between MHC-β loss and pathogenicity for certain *MYH7* variants. To further support this novel finding, we leveraged an observation that hiPSC-CMs within a single genotype and differentiation exhibit variable mTagBFP2 depletion (**Fig. 3A**) to perform comparative analyses of FSCA (a proxy for cell size) in mTagBFP2^low^ versus mTagBFP2^high^ hiPSC-CMs using flow cytometry (**Fig. 3E**). These analyses revealed a significant increase in FSCA in mTagBFP2 low versus high MYH7^WT-eGFP/S866F-mTagBFP2^ and MYH7^WT-eGFP/S866W -mTagBFP2^ hiPSC-CMs (average 13% and 12%, respectively) relative to WT hiPSC-CMs (**Fig. 3E**). Consistent with our previous study [19], we do not observe a significant difference in FSCA in mTagBFP2 low versus high MYH7^WT-eGFP/M852T-mTagBFP2^ hiPSC-CMs relative to WT hiPSC-CMs (**Fig. 3E**), suggesting cellular hypertrophy correlates with the severity of MHC-β loss. Taken together, these clonal analyses validate our VAMP-seq results, indicating multiplexed FACS-based MHC-β abundance phenotyping is an accurate approach to assess *MYH7* variant effect at scale.

### A multiplexed assessment of survival in hiPSC-CMs containing a library of heterozygous *MYH7* missense variants accurately predicts pathogenicity

Previous reports by our group [27] and others [11] demonstrated hiPSC-CMs with pathogenic *MYH7* missense variants (MYH7^E848G/+^ or MYH7^R723C/R723C^, respectively) have progressively reduced numbers of hiPSC-CMs relative to isogenic controls, thus, we hypothesized that a multiplexed survival assay would segregate hiPSC-CMs with P/LP *MYH7* variants from hiPSC-CMs with Syn *MYH7* variants. To this end, we examined the final frequencies of five P/LP variants and three Syn variants in hiPSC-CMs at T25 and T47 relative to their initial frequencies at the hiPSC stage, when *MYH7* is not expressed/functional [30] (hiPSC and T25 hiPSC-CM NGS data from Friedman *et al*. [19]) (**Fig. 4A**; **Supp. Fig. S5A**; **Supp. Tables S1-3**). This analysis revealed that at T25, 5/5 P/LP variants were less frequent than 3/3 Syn variants (average 50% reduction) (**Supp. Fig. S5A**) and that at T47, 5/5 P/LP variants were less frequent than 3/3 Syn variants (average 59% reduction) (**Fig. 4A**). Note that two rare P/LP variants (p.Met852Arg and p.Lys865Arg) did not reach NGS frequency thresholds at T47 for quantification, and were thus considered ‘Lost’.

**Fig. 4.**
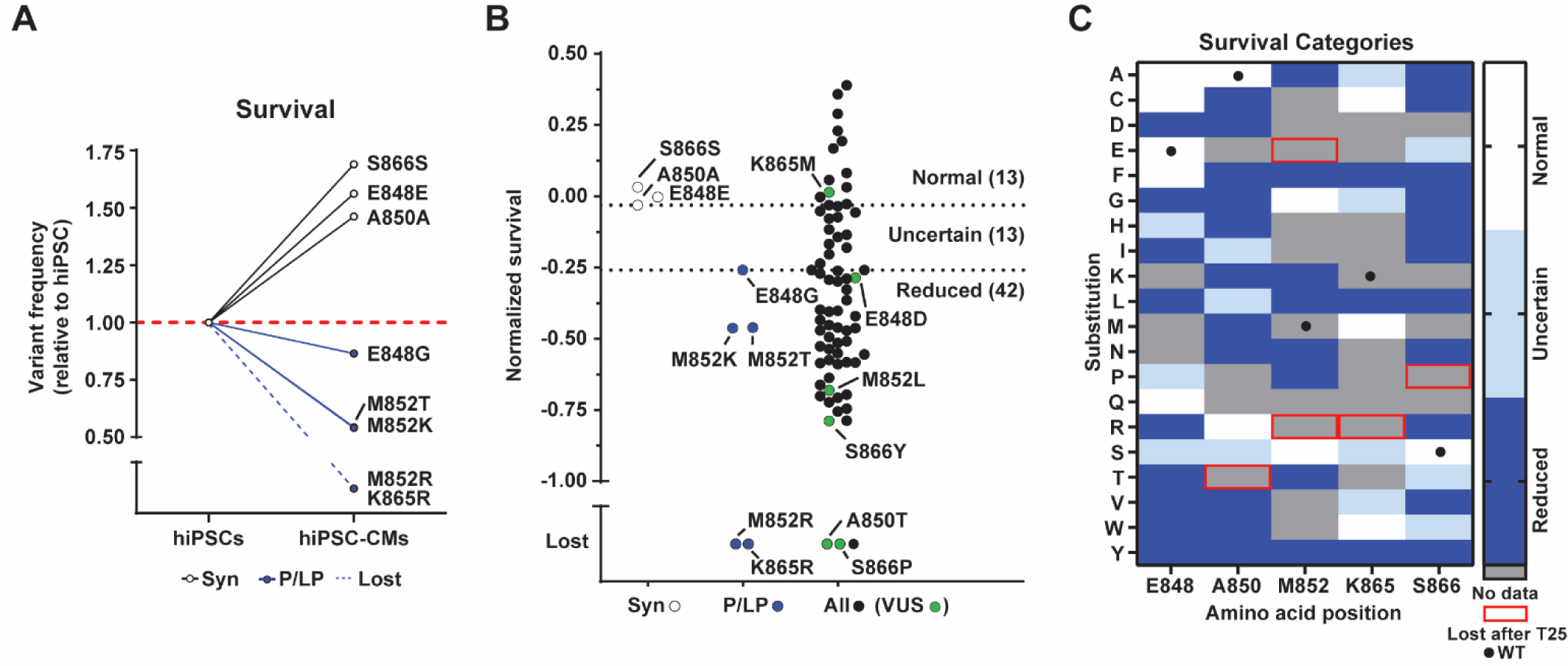
Survival in a library of heterozygous *MYH7* missense variant hiPSC-CMs. **A** Variant frequency of MYH7^WT-eGFP/Variant-mTagBFP2^ hiPSC-CMs with single synonymous (Syn; white) or pathogenic/likely pathogenic (P/LP; blue) *MYH7* variants assessed at T47 (*n* = 2 independent replicates) relative to starting frequencies as hiPSCs (*n* = 1 replicate) (hiPSC dataset from Friedman *et al.* [19]). T = days after the activation of differentiation. **B** Mean normalized T47 survival scores for synonymous (Syn; white), pathogenic/likely pathogenic (P/LP; blue), and all variants (black) (*n* = 2 independent replicates). Variants of unknown significance (VUS) scores in ‘All’ column = green dots. Upper dotted line = lowest ‘Normal’ survival score; lower dotted line = highest ‘Reduced’ survival score; scores between ‘Normal’ and ‘Reduced’ are ‘Uncertain’. **C** Position map displaying T47 normalized categorical survival scores (relative to hiPSCs) for each mutagenized residue (vertical) relative to amino acid substitutions (horizontal). Grey = no data. Red box = present in T25 hiPSC-CMs and hiPSCs but ‘Lost’ at T47. Black dots = WT residues. Dashed red line in **A** indicates normalized starting variant frequencies in hiPSCs. See **Supplemental Figure S5** for T25 and T47 *MYH7* variant hiPSC-CM survival data and T47 replicate data.

We first calculated survival scores across all 68 amino acid substitutions present in T47 hiPSC-CMs relative to hiPSCs (normalized to Syn variants) and found scores between independent replicates were highly consistent (R^2^ = 0.8404), indicating assessment of multiplexed survival detects a reproducible phenotype (**Supp. Fig. S5D**). 13/68 (19.1%) amino acid substitutions resulted in survival greater than the lowest Syn variant and were considered functionally Normal (**Fig. 4B-C**; **Supp. Fig. S5E**). 42/68 (61.8%) substitutions resulted in survival lower than the highest P/LP variant and were considered functionally Reduced, including 3/4 VUS (**Fig. 4B-C**; **Supp. Fig. S5E**). Interestingly, the p.Lys865Met VUS has Normal survival yet Reduced MHC-β abundance, suggesting potential dysfunction in this case could be caused by MHC-β loss alone (**Fig. 2-4**). Note five variants (p.Ala850Thr, p.Met852Glu, and p.Met852Arg; p.Lys865Arg; p.Ser866Pro) were so rare in T47 hiPSC-CMs that they failed to pass NGS thresholds and their frequencies were not quantifiable at T47. Moreover, of these five ‘lost’ variants, two are P/LP (p.Met852Arg and p.Lys865Arg) and two are VUS (p.Ala850Thr and p.Ser866Pro), further correlating variant loss with clinical pathogenicity. 13/68 (19.1%) amino acid substitutions resulted in survival lower than the lowest Syn variant yet higher than the highest P/LP variant and were considered functionally Uncertain (**Fig. 4B-C**; **Supp. Fig. S5E**).

We also calculated survival scores across all 72 amino acid substitutions present in T25 hiPSC-CMs relative to hiPSCs (normalized to Syn variants) and found that 16/72 (22.2%) amino acid substitutions resulted in survival greater than the lowest Syn variant and were considered functionally Normal (**Supp. Fig. S4B-C**). The remaining 60/72 (87.8%) substitutions resulted in survival lower than the highest P/LP variant and were considered functionally Reduced, including all six VUS (**Supp. Fig. S4B-C**). Note six unlabeled variants (p.Ala850Pro, p.Ala850Gln, p.Lys865Asp, p.Ser866His, p.Ser866Met, and p.Ser866Gln) were so rare in T25 hiPSC-CMs, that they failed to pass NGS thresholds and their frequencies were not quantifiable at T25 (**Supp. Fig. S4B-C**). Taken together, this multiplexed hiPSC-CM survival assay segregates all tested P/LP variants (T25 5/5; T47 5/5) and most VUS (T25 5/6; T47 3/4) from Syn variants, indicating this is an additional, accurate approach to assess *MYH7* variant effect at scale.

### MHC-β abundance and survival phenotyping identifies shared and unique variant effects

Leveraging multiple independent, multiplexed functional assays provides greater confidence for variant effect calling of variants with consistent scores across both assays and provides an opportunity to distinguish between different mechanisms of pathobiology. To this end, we compared normalized MHC-β abundance scores relative to corresponding normalized survival scores for all 68 *MYH7* amino acid substitutions with replicate data (**Fig. 5**). This analysis revealed 33/68 (48.5%) agreement across functional categories between assays, with 23/33 (69.7%) consensus substitutions called functionally Abnormal in both assays, 7/33 (21.2%) consensus substitutions called functionally Normal in both assays, and 3/33 (9.1%) consensus substitutions called functionally Uncertain in both assays (**Fig. 5**). 3/4 VUS substitutions were considered functionally Abnormal in both assays, while 1/4 VUS (p.Lys865Met) was considered functionally Abnormal only in the MHC-β abundance assay; three other substitutions (p.Glu848Gln, p.Met852Gly, and p.Met852Ser) also had similar conflicting results. Together, those four substitutions and fourteen others considered functionally Normal in the MHC-β abundance assay, yet functionally Abnormal in the survival assay, have conflicting effects accounting for 18/68 (26.5%) of substitutions and likely represent cases where a unique pathogenic mechanism is dominant. The remaining 20/68 (29.4%) substitutions were categorized as functionally Uncertain in one or more assays and will require further examination of additional phenotypes to determine variant effect.

**Fig. 5.**
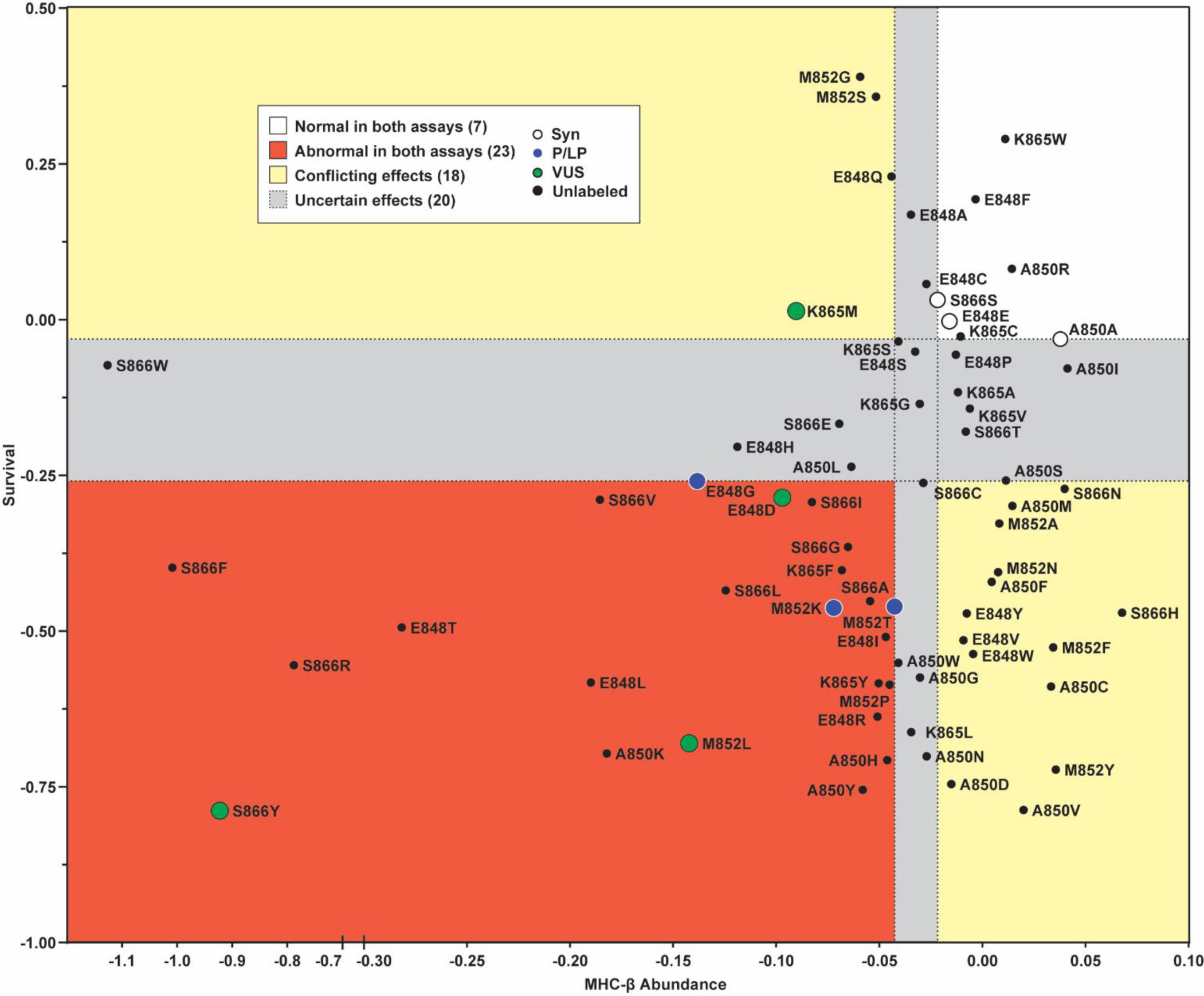
Comparison of MHC-β abundance to survival across a library of heterozygous *MYH7* missense variant hiPSC-CMs. Plot displaying mean normalized MHC-β-mTagBFP2 abundance scores (x-axis) relative to mean normalized hiPSC-CM survival scores (T47; y-axis) for synonymous (Syn; white), pathogenic/likely pathogenic (P/LP; blue), variants of unknown significance (VUS; green), and unlabeled variants (black) (*n* = 2 independent replicates). The white domain indicates ‘Normal’ variant effect in both assays, the orange domain indicates ‘Abnormal’ variant effects in both assays, the yellow domains indicate ‘Conflicting effects’ between assays, and the grey domain indicates ‘Uncertain effects’ in at least one assay.

Four amino acid substitutions encoded for by VUS or variants with conflicting interpretations (considered VUS in this study) (p.Glu848Asp, p.Met852Leu, p.Lys865Met, and p.Ser866Tyr) all resulted in reduced MHC-β abundance and survival, save p.Lys865Met (**Fig. 2** and **4-5**). Patients with these *MYH7* variants have disease profiles including HCM, cardiomyopathy, and/or cardiovascular phenotypes [2]. The multiplexed data generated here indicate p.Glu848Asp, p.Met852Leu, and p.Ser866Tyr are functionally abnormal and should likely be classified as pathogenic (P). In addition, 17 unlabeled missense variants have both Abnormal MHC-β abundance and survival (**Fig. 5**) and should likely be classified as P/LP if patients with those variants are identified in the future. However, we acknowledge this study lacks sufficient controls to make conclusive classifications until more definitive evidence is generated.

In summary, this study demonstrates the synergy of hiPSC differentiation capacity into disease-relevant cell types combined with multiplexed phenotyping assays to enable a proof-of-concept DMS of *MYH7*, a clinically-actionable gene, in stem cell derivatives for the first time. These experiments reveal that multiplexed MHC-β abundance and cardiomyocyte survival assays accurately predict pathogenicity of all tested P/LP *MYH7* missense variants, highlighting that concordant predictions from these independent assays increase the confidence of variant effect calling. This work highlights exciting new opportunities to study variant effect in more clinically-actionable genes and disease-relevant cellular contexts.

## Discussion

Here we report, to our knowledge, the first DMS executed in a human induced pluripotent stem cell (hiPSC) derivative. hiPSCs have the capacity to differentiate to nearly any cell type in the body and numerous protocols enable directed differentiation of hiPSCs to specific cell types [8, 9]. Importantly, hiPSC derivatives have been successfully utilized to investigate the pathogenicity of missense variants in numerous genes because they can manifest salient disease phenotypes [10, 27, 28, 31, 32]. We reasoned that mutagenesis in hiPSCs followed by differentiation would enable DMS of previously inaccessible, clinically-actionable genes in nearly any disease-relevant cell type. As a proof of concept, we leveraged gene-edited hiPSCs that our group previously generated targeting *MYH7* [19], which encodes a sarcomeric thick filament protein (MHC-β) expressed in cardiac and skeletal myocytes. These isogenic hiPSCs (derived from a healthy Japanese male background (WTC11; XY)) contain a biallelically-edited heterozygous *MYH7* missense library consisting of five amino acid positions (848, 850, 852, 865, and 866) mutagenized to saturation within a *MYH7* cDNA transgene tagged with mTagBFP2, while the other *MYH7* allele contains WT *MYH7* cDNA tagged with eGFP [19]. A limitation of this intronless, cDNA-based gene editing approach is that intronic variants and variants effecting intron/exon splicing cannot be examined, however, these types of variants account for <2.5% of *MYH7* missense variants reported in ClinVar [2]. The mutagenized residues are in the proximal S2 domain, a hotspot for mutations that cause HCM and DCM [29]. Directed differentiation of this *MYH7* variant hiPSC library to cardiomyocytes provides the correct cellular context to study *MYH7* variants associated with cardiomyopathies – cancer cell lines traditionally used for DMS express neither *MYH7* nor the milieu of cardiomyocyte-specific chaperones that regulate MHC-β expression and function.

We first sought to identify a multiplexable phenotype that accurately segregates an HCM-causing *MYH7* mutant, MYH7^E848G/+^ [10], from WT cells. Examination of MHC-β abundance in failing and non-failing patient heart samples and in isogenic, hiPSC-CMs with the p.Glu848Gly P/LP mutation showed reduced MHC-β abundance in cardiomyocytes. We next sought to determine the utility of the MHC-β depletion phenotype for multiplexable assessment of variant effect in the *MYH7* variant hiPSC-CM library at scale. We leveraged both *MYH7* alleles being tagged with different fluorescent proteins (a design that enabled CRaTER enrichment [19]) to perform MHC-β-based variant abundance by massively parallel sequencing (VAMP-seq). These data revealed that non-synonymous P/LP variants consistently express less MHC-β than synonymous (Syn) variants. While *MYH7* nonsense variants are generally well-tolerated [3, 33], we posit that *MYH7* missense variants that reduce MHC-β abundance could cause pathogenicity through an independent and/or indirect mechanism to haploinsufficiency.

Interestingly, we found certain substitutions of the p.Ser866 residue cause the most severe MHC-β depletions. We hypothesize these substitutions could (1) eliminate the p.Ser866 phosphorylation site in MHC-β [34] and interfere with the function of that posttranslational modification, and/or (2) sterically block MHC-β homodimerization by disrupting the hydrogen bond between the p.Ser866 residue of one monomer and the p.Glu867 residue of the other [35]. We found that 4/4 variants of unknown significance (VUS) also have reduced MHC-β abundance relative to controls, providing functional evidence that the cardiac phenotype(s) observed in patients with these VUS could be caused by reduced MHC-β abundance. Future MAVEs in extended regions of *MYH7* will aid in determining if MHC-β loss can accurately predict pathogenicity across all *MYH7* domains or if this phenotype is restricted to the proximal S2 domain. Importantly, VAMP-seq does not directly measure other aspects of MHC-β function (e.g., ATP hydrolysis or MyBP-C binding), therefore it is possible that variants in these MHC-β functional domains could cause no change in MHC-β abundance and yet remain pathogenic. We suggest that conclusions about benignness from our VAMP-seq results should be avoided.

In addition to multiplexed assessment of MHC-β abundance, we assessed survival of *MYH7* variant hiPSC-CMs in a multiplexed fashion. This functional assay compares variant frequencies in cardiomyocytes relative to their frequency in hiPSCs, when *MYH7* is not expressed [30] and should not exert any functional effect. In line with previous examinations of single, pathogenic *MYH7* missense variants [11, 27], we demonstrate that multiplexed assessment of survival correctly segregates all tested P/LP variants and 3/4 VUS from Syn variants. HCM patients with pathogenic *MYH7* variants can develop enhanced cardiac fibrosis by an unknown mechanism [29]; our data suggests that cardiomyocyte loss could play a role in the development of cardiac fibrosis by skewing cellular proportions toward fibroblasts (which do not express *MYH7*). Importantly, nearly 50% of functional scores from this independent functional assay are consistent with MHC-β abundance functional scores, providing a high level of confidence for those variants. Interestingly, pathogenic variants that consistently scored as functionally Abnormal across the MHC-β abundance and survival assays have inconsistent consequences on contractile function. For example, we have shown that p.Met852Thr variant hiPSC-CMs are hypercontractile [19], while p.Glu848Gly variant hiPSC-CMs are hypocontractile [10, 27]. These data suggest that while Abnormal MHC-β abundance and survival scoring cannot predict the direction of consequences on force generation, it can predict contractile dysfunction generally.

As cardiomyocytes function in a syncytium *in vivo*, it was initially uncertain whether this ‘cohort in a dish’ of different *MYH7* variant hiPSC-CMs could be co-cultured during differentiation and during multiplexed assessment(s) of variant effect without phenotypic interference via paracrine signaling. However, given that *MYH7* encodes an intracellular filament protein, we expected there would be few intercellular effects on MHC-β abundance. Indeed, examination of a subset of *MYH7* variant hiPSC-CMs individually differentiated and examined for MHC-β abundance revealed a high correlation to the multiplexed VAMP-seq analyses, suggesting that pooled variant library co-culture and phenotyping are subject to minimal paracrine effects.

It remains unclear whether the MHC-β loss phenotype described here is disease-causing or is a downstream readout of a pathogenic mechanism. Interestingly, within a single genotype and differentiation, one set of hiPSC-CMs decrease mTagBFP2 abundance while another set appears to have WT-like mTagBFP2 abundance, suggesting MHC-β loss is partially stochastic; this variability in MHC-β loss is also observed when sampling MHC-β expression in different sections of the human heart. Understanding the factors that determine MHC-β loss is key for maximizing the dynamic range of MHC-β abundance in future VAMP-seq experiments. Preliminary evidence indicates that severe MHC-β loss is associated with increased cardiomyocyte size (a common HCM phenotype [28]) suggesting that MHC-β loss is potentially deleterious, however, the direction of causation remains unclear. MHC-β loss and hiPSC-CM survival should be further leveraged as means to identify novel multiplexable markers of pathogenicity that can be utilized in future *MYH7* DMS. Broadly, understanding the relationship between MHC-β loss, reduced hiPSC-CM survival, and cardiomyopathies will have important implications for the development of therapeutic strategies for patients with *MYH7* mutations.

## Conclusions

To date, DMS have primarily targeted genes with salient phenotypes in cancer cell lines. Studying variant effect of clinically-actionable genes in disease-relevant contexts has been hindered by low gene-editing efficiency of relevant cell types. Building from our previous work showing CRaTER enrichment of gene-edited cells enables the generation of hiPSC variant libraries, we leveraged the differentiation capacity of hiPSCs to differentiate a *MYH7* missense variant library to cardiomyocytes, the effected cell type in *MYH7*-associated cardiomyopathies. Assessment of *MYH7* variant effect in hiPSC-CMs by VAMP-seq and hiPSC-CM survival accurately segregated all tested synonymous controls from pathogenic variants and the majority of VUS. This study bridges DMS with the differentiation power of hiPSCs to enable variant effect studies in new frontiers. Together, the connection of these technologies expands capacities to classify variants in previously inaccessible, clinically-actionable genes and presents opportunities to discover novel mechanisms of pathophysiology at scale.

## Methods

No animals were used in this study.

### Human heart sample isolation

Immediately upon explant of the *MYH7* (p.Glu848Gly) failing heart, random samples were obtained from the septum and the left ventricular free wall. The samples were all immediately flash-frozen in liquid nitrogen and then stored long-term in -80° C.

### Western blots

#### Lysate preparation

Human cardiac biopsy tissue was obtained with appropriate consent. Tissues were minced, homogenized, and lysed in Pierce RIPA lysis buffer (Thermo Fisher; cat. no. 89901) supplemented with Halt protease and phosphatase inhibitor (Invitrogen; cat. no. 78443) and dithiothreitol (Roche; cat. no. 3483-12-3). Lysates were rocked at 4° C for 20 min and centrifuged for 10 min at 15,000 × *g*, with super-natant collected. Total protein concentration of supernatant was assessed through a Bradford assay (Bio Rad; cat. no. 5000006) with 560 nm absorbance per manufacturer’s protocol using BSA standards (Thermo Fisher; cat. no. 23208). Lysates were diluted with 4X Laemmli SDS Sample Buffer (Bio Rad; cat. no. 1610747) to 1 μg/μL.

#### Electrophoresis and Staining

30 μg of lysate was loaded per lane in Mini Protean 4–15% polyacrylamide gels (Bio Rad; cat. no. 4508084). Electrophoresis was run at 120 V for 50 min in 1X tris-glycine-SDS running buffer (25 mM tris base (Sigma; cat. no. T1503-1KG), 190 mM glycine (Fisher; cat. no. BP381-1), and 0.1% sodium dodecyl sulfate (Fisher; cat. no. BP243-1)). Proteins were transferred to Immobilon-P membranes (Millipore; cat. no. IPVH85R) at 120 V for 50 min on ice in 1X tris-glycine-methanol transfer buffer (25 mM tris base, 190 mM glycine, and 20% methanol (Fisher; cat. no. A412P-4)). Membranes were blocked with 5% milk (Bio Rad; cat. no. 1706404) in 1X tris buffer saline with Tween-20 (TBST; 20 mM tris base and 150 mM Tween-20 (Sigma; cat. no. P9416-100mL)) for 120 min at 22° C. Membranes were washed three times with 1X PBS and incubated with appropriate primary antibody (see table below) diluted in 4% BSA (Sigma; at. No. A9418-50G) in 1X PBS overnight at 4° C. Membranes were washed three times with 1X PBS and incubated with appropriate secondary antibody diluted in 1% BSA in 1X TBST for 60 min at 22° C. Membranes were washed three times with 1X PBS and incubated with Clarity Max ECL substrate (Bio Rad; cat. no. 1705061) for 3 min. Chemiluminescence images were obtained using the ChemiDoc imaging system (Bio Rad; cat. no. 17001401). Volumetric band intensities were analyzed using Bio Rad Image Lab software (Version 6.1.0, Bio Rad).

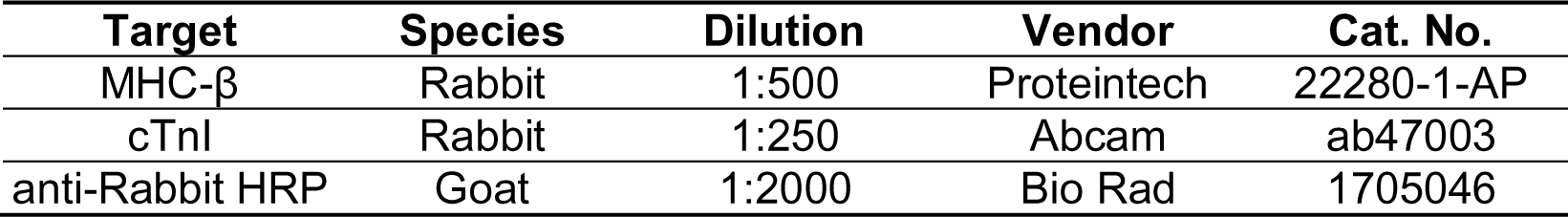

| Target | Species | Dilution | Vendor | Cat. No. |
| --- | --- | --- | --- | --- |
| MHC-β | Rabbit | 1:500 | Proteintech | 22280-1-AP |
| cTnl | Rabbit | 1:250 | Abcam | ab47003 |
| anti-Rabbit HRP | Goat | 1:2000 | Bio Rad | 1705046 |

### Stem cell culture and maintenance

All cells maintained at 37° C in 5% CO_2_ as previously described in Friedman et al., 2023. Human induced pluripotent stem cells (hiPSCs), including wildtype (WT) WTC 11 hiPSCs (XY; a gift from Bruce Conklin) and subclones derived from these cells were culture on Matrigel (Corning; cat. no. 354277) and fed every other day with stem cell media: mTeSR Plus with supplement (STEMCELL Technologies; cat. no. 100-0276) and 0.5% penicillin-streptomycin (ThermoFisher; cat. no. 15140122). When 80% confluent (approximately every 4 days) hiPSCs were dissociated using 0.5 mM EDTA (Invitrogen; cat. no. 15575-038) in DPBS (Invitrogen; cat. no. 14190250) and resuspended in stem cell media supplemented with 10 µM Y-27632 dihydrochloride (Rho kinase [ROCK] inhibitor; Tocris; cat. no. 1254). hiPSCs were used for ≤ 10 passages after thawing.

### Cardiac-directed differentiation and culture

Small molecule cardiac-directed differentiation was performed following previous protocols [19]. On differentiation day T-1, hiPSCs were dissociated as above and replated into a 24 well-plate at 1.32 × 10^5^ cells/cm^2^. On T0 (activation day), the stem cell media with ROCK inhibitor was removed, cells washed once with DPBS, and fed with RBA media: RPMI (Invitrogen; cat. no. 11875135), 0.5 mg/ml bovine serum albumin (Sigma; cat. no. A9418), 0.213 mg/ml ascorbic acid (Sigma; cat. no. A8960), and 0.5% penicillin-streptomycin supplemented with 4 µM CHIR-99021 (Cayman; cat. no. 13122). On T3, CHIR-99021-containing media was removed, cells washed once with DPBS, and fed with RBA supplemented with 2 µM Wnt-C59 (Selleck; cat. no. S7037). On T5, Wnt-C59-containing media was replaced with RBA. On T7 (and every other day afterwards), media was replaced with cardiomyocyte media: RPMI, B27 plus insulin (Invitrogen; cat. no. 17504044), and 0.5% penicillin-streptomycin.

To enrich for cardiomyocytes using lactate selection, T19-21 cells were washed once with DPBS and then dissociated using 37° C 0.5 mM EDTA with 0.5% trypsin (Invitrogen; cat. no.15090046) in DPBS for 7 min at 37° C in 5% CO_2_. The cells were collected, 25 µU DNAse I (Sigma; cat. no. 260913) was added, and then cells were triturated using a serological pipette until homogenized. The reaction was stopped using an equal volume of cardiomyocyte media supplemented with 5% fetal bovine serum (FBS; Biowest: cat. no. S1520) and filtered using a 100 µm strainer. Cells were centrifuged at 300 × *g* for 3 min and the supernatant aspirated then the pellet was resuspended in cardiomyocyte media with 5% FBS and replated at 5.3-8.8 × 10^5^ cells/cm^2^ in 10-cm plates and incubated overnight at 37° C in 5% CO_2_. For the next 4 days, cells were fed daily with lactate media: DMEM (Invitrogen; cat. no. A1443001), 2 mM ʟ-glutamine (Invitrogen; cat. no. 25030081), 0.5% penicillin-streptomycin, and 4 mM sodium ʟ-lactate (Sigma; cat. no. 71718).

Following lactate selection, hiPSC-CMs were washed once with DPBS and then dissociated using 37° C 0.5 mM EDTA in PBS and 0.5% trypsin for 5 min at 37° C in 5% CO_2_. Cells were collected and triturated using a serological pipette until homogenized. The reaction was stopped with an equal volume of DMEM cardio media (DMEM (Invitrogen; cat. no. 10313-021), 0.5% penicillin-streptomycin, 2 mM ʟ-glutamine, and B27 plus insulin) supplemented with 5% FBS and 25 µU DNAse I. Cells were centrifuged at 300 × *g* for 3 min and the supernatant aspirated then the pellet was resuspended in DMEM cardio media with 5% FBS and replated at 6.6 × 10^4^ cells/cm^2^ in 24-well plate wells and incubated overnight at 37° C in 5% CO_2_. The following day, and every other day thereafter, cells were fed with DMEM cardio media.

### Flow cytometry

hiPSC-CMs were harvested for flow cytometry as above and filtered using a 100 µm strainer prior to centrifugation at 300 × *g* for 3 min. The supernatant was aspirated, and the pellet was resuspended in 300 µL PBS with 5% FBS for flow cytometry using a CantoII (speed high; BD).

### Fluorescence-activated cell sorting (FACS)

hiPSC-CMs were harvested as above but instead resuspended in 100-300 µL FACS sorting buffer: DPBS, 1% penicillin streptomycin, 2% FBS, 10 mM HEPES (Invitrogen; cat. no. 15630-080), and 25 µU DNAse I and sorted using a FACS AriaIII Cell Sorter (sort speed 1; BD).

### Genomic DNA isolation

Genomic DNA (gDNA) was isolated using the DNeasy Blood & Tissue Kit following manufacturer’s instructions (Qiagen; cat. no. 69504).

### Next generation sequencing (NGS)

#### hiPSC-CM gDNA amplicon preparation

Following genomic DNA isolation using the DNeasy Blood & Tissue Kit following manufacturer’s instructions (Qiagen; cat. no. 69504), amplicons were prepared as described in Friedman *et al*. [19].

#### Sequencing parameters

Sequencing was performed on a MiSeq instrument using a MiSeq Reagent Kit v3 (150-cycle; Illumina; cat. no. MS-102-3001) following standard procedures. Custom index and read primers were loaded to sequence the amplicons, with 8 cycles allocated for each index and 60 cycles for each read 1 and read 2 sequences. To increase sequence diversity and improve base calling, 10-20% PhiX control DNA (Illumina; cat. no. FC-110-3001) was spiked into the pooled 2 nM DNA amplicon library. To optimize clustering and read output, kits were loaded with a total of 22 pmol of the DNA amplicon library.

#### Data analysis

After demultiplexing on index reads using bcl2fastq version 2.20, paired-end 57-bp reads were merged into a q30 consensus using Paired-End reAd mergeR 0.9.11 and counted [36]. T47-48 *MYH7* variant library cardiomyocytes variants passed threshold if they were present in the hiPSC library (after hiPSC-specific thresholding) [19]. To increase stringency, the least frequent 13% of variants were excluded. Cardiomyocyte variant frequency was calculated as (variant read count)/(total variant read counts) with only NNK variant codons (besides the p.E848= (GAA) spike-in considered. Read counts for codons encoding the same residue were averaged to generate a final variant count.

### Variant abundance by massively parallel sequencing (VAMP-seq)

T47-48 *MYH7* variant library cardiomyocytes were harvested for sorting as above. GFP^+^ cells were sorted into three bins based on variable mTagBFP2 positivity: low (bottom ∼10%), mid (middle ∼45%), and high (top ∼45%). The geometric mean of mTagBFP2 intensity for each bin (I) was calculated using FlowJo (v10; BD) and the number of sorted cells was used to quantify the sorting bin frequencies (B) calculated as (bin cell count)/(total sorted cell count across bins). The frequency of each variant (F) within each bin was calculated as (final variant count)/(total final variant counts). Abundance scores were weighted by bin-specific mTagBFP2 intensities and sorting bin frequencies and then log transformed as below:

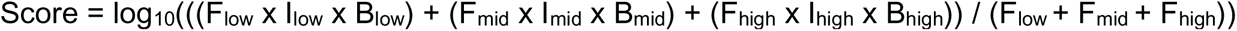

Abundance scores were normalized to synonymous variant scores (Syn; p.E848=, p.A850=, and pS866=) by subtracting the average synonymous score from each variant’s score. Variants with normalized negative scores have reduced abundance relative to Syn variants and vice versa.

### Cardiomyocyte survival analyses

Initial variant frequencies in hiPSCs were calculated using NGS data from Friedman *et al*. [19]. Read counts for codons encoding the same residue were averaged to generate a final variant count. The percentage of each variant was calculated as ((100)(final variant count))/(total final variant counts).

#### T25 hiPSC-CM analysis

T25 hiPSC-CM variant percentages were calculated the same as with hiPSCs above and also used data from Friedman *et al*. [19]. The fold change of T25 hiPSC-CM variant percentages relative to corresponding hiPSC percentages was calculated as (T25 hiPSC-CM variant percentages)/(hiPSC variant percentage). This fold change was normalized to synonymous variant fold changes (Syn; p.E848=, p.A850=, and pS866=) by subtracting the average synonymous fold change from each variant’s fold change and then performing a log_10_ transformation. Variants with normalized negative scores have reduced survival at T25 relative to Syn variants and vice versa.

#### T47-48 hiPSC-CM analysis

The T47-48 hiPSC-CM survival analysis used the same NGS data generated here and used for VAMP-seq. T47-48 hiPSC-CM variant percentages were calculated on a bin by bin basis. These bin-specific variant percentages were then weighted to account for each sorting bin frequency (B) and calculated as (bin percentage)(B) and then summed to reconstruct a variant percentage. The fold change of T47 hiPSC-CM variant percentages relative to corresponding hiPSC percentages was calculated as (T47 hiPSC-CM variant percentages)/(hiPSC variant percentage). This fold change was normalized to synonymous variant fold changes (Syn; p.E848=, p.A850=, and pS866=) by subtracting the average synonymous fold change from each variant’s fold change and then performing a log_10_ transformation. Variants with normalized negative scores have reduced survival at T47 relative to Syn variants and vice versa.

### Confocal microscopy

Spinning disk microscopy was performed on a Nikon Eclipse Ti equipped with a Yokogawa CSU-W1 spinning disk head, Andor iXon LifeEMCCD camera, and 60x Plan Apo objective Images were taken at 37° C in 5% CO_2_.

### Statistical analysis

Bar graphs shown mean and error bars indicate standard error of the mean. Paired and unpaired *t*-tests, as well as ordinary and repeat measure one-way ANOVA were used to determine statistical significances as shown above bars (*p* < 0.05; *p* > 0.05 considered not significant (NS)).

## Abbreviations

DMS: deep mutational scan
FACS: fluorescence-activated cell sorting
MAVE: multiplexed assay of variant effect
MHC-β: myosin heavy chain-β
MYH7: myosin heavy chain 7
NGS: next generation sequencing
VAMP-seq: variant abundance by massively parallel sequencing

## Declarations

### Ethics approval and consent to participate

The study was conducted in compliance with the requirements stipulated in the U.S. Department of Health and Human Services (DHHS) Protection of Human Subjects regulations at 45 CFR 46, and approved by the Institutional Review Boards of Seattle Children’s Hospital (FWA# 00002443; IRB protocol no. 12856) and the University of Washington (FWA# 00006878; IRB protocol numbers 1553 and 35358). Human subjects gave informed consent for this research study of their specimens and medical information.

## Consent for publication

Not applicable.

## Availability of data and materials

All data are available in the main text or the supplementary materials. The GEO submission number associated with the raw data presented in this manuscript will be provided. Human data presented in this study are openly available at Zenodo (doi: 10.5281/zenodo.7563492).

## Authors’ contributions

Conceptualization, C.E.F., S.F., S.P., L.M.S., D.M.F., and K-C.Y.; Methodology, C.E.F., S.F., and S.P.; Software, S.P.; Validation, C.E.F.; Formal Analysis, C.E.F.; Investigation, C.E.F., S.F., S.P., W-M.C., L.T., L.S.C., A.M., D.A., A.L., and S.D.F.; Resources, L.M.S. and D.M.F.; Data Curation, C.E.F.; Writing – Original Draft, C.E.F.; Writing – Review & Editing, C.E.F., S.F., S.P., W-M.C., A.L., C.E.M., L.M.S., D.M.F., and K-C.Y.; Visualization, C.E.F.; Supervision, C.E.F., A.S.-O., and K-C.Y.; Project Administration, K-C.Y.; Funding Acquisition, C.E.F., S.F., S.P., A.L., C.E.M., L.M.S., D.M.F., and K-C.Y.

## Acknowledgements

We are thankful to the staff of the Cell Analysis Facility at the University of Washington and the Mike and Lynn Garvey Cell Imaging Core at the Institute for Stem Cell and Regenerative Medicine (University of Washington) and its director, Dale Hailey.

## Supplemental Figures and Tables

**Table of Contents**

Supplemental Figure S1 Complete stain-free and Western blots, associated with Fig. 1.

Supplemental Figure S2 Pooled analyses of *MYH7* variant hiPSC-CM library mTagBFP2 intensity and FSCA, associated with Fig. 2.

Supplemental Figure S3 VAMP-seq replicate data, associated with Fig. 2.

Supplemental Figure S4 Confocal microscopy of clonal heterozygous *MYH7* variant hiPSC-CMs, related to Fig. 3.

Supplemental Figure S5 T25 *MYH7* variant hiPSC-CM survival and T47 *MYH7* variant hiPSC-CMs replicate survival data, associated with Fig. 4.

Supplemental Table S1 T47 hiPSC-CM DNA variant library NGS data, VAMP-seq analyses, and survival analyses.

Supplemental Table S2 T25 hiPSC-CM DNA variant library NGS data and survival analysis.

Supplemental Table S3 hiPSC DNA variant library NGS data and survival analysis.

**Supplemental Figure S1.**
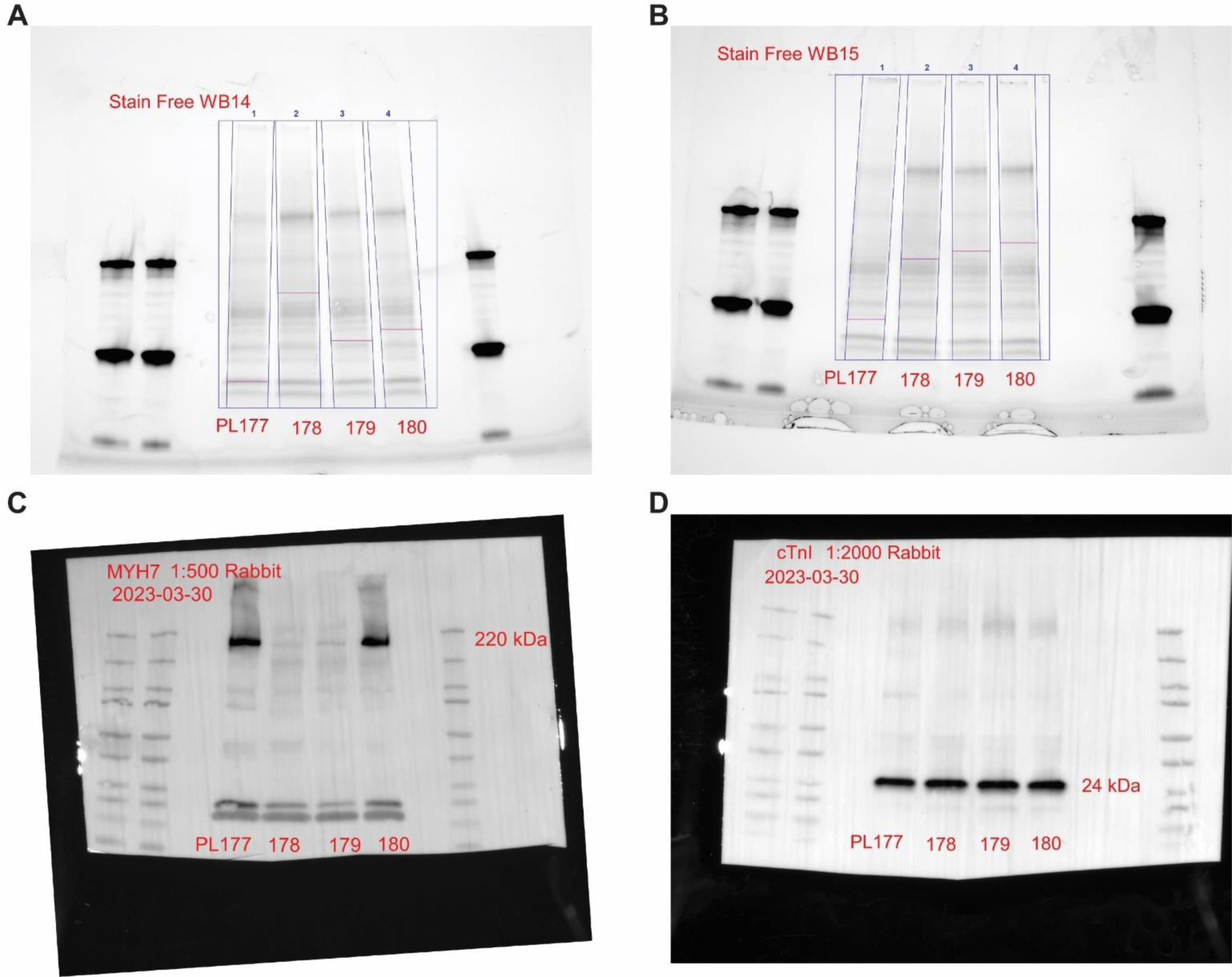
Complete stain-free and Western blots, associated with Fig. 1. Complete stain-free blots used to normalize protein loading for each lane of the MHC-β (**A**) and cTnI (**B**) blots. Complete MHC-β (**C**) and cTnI (**D**) Western blots.

**Supplemental Figure S2.**
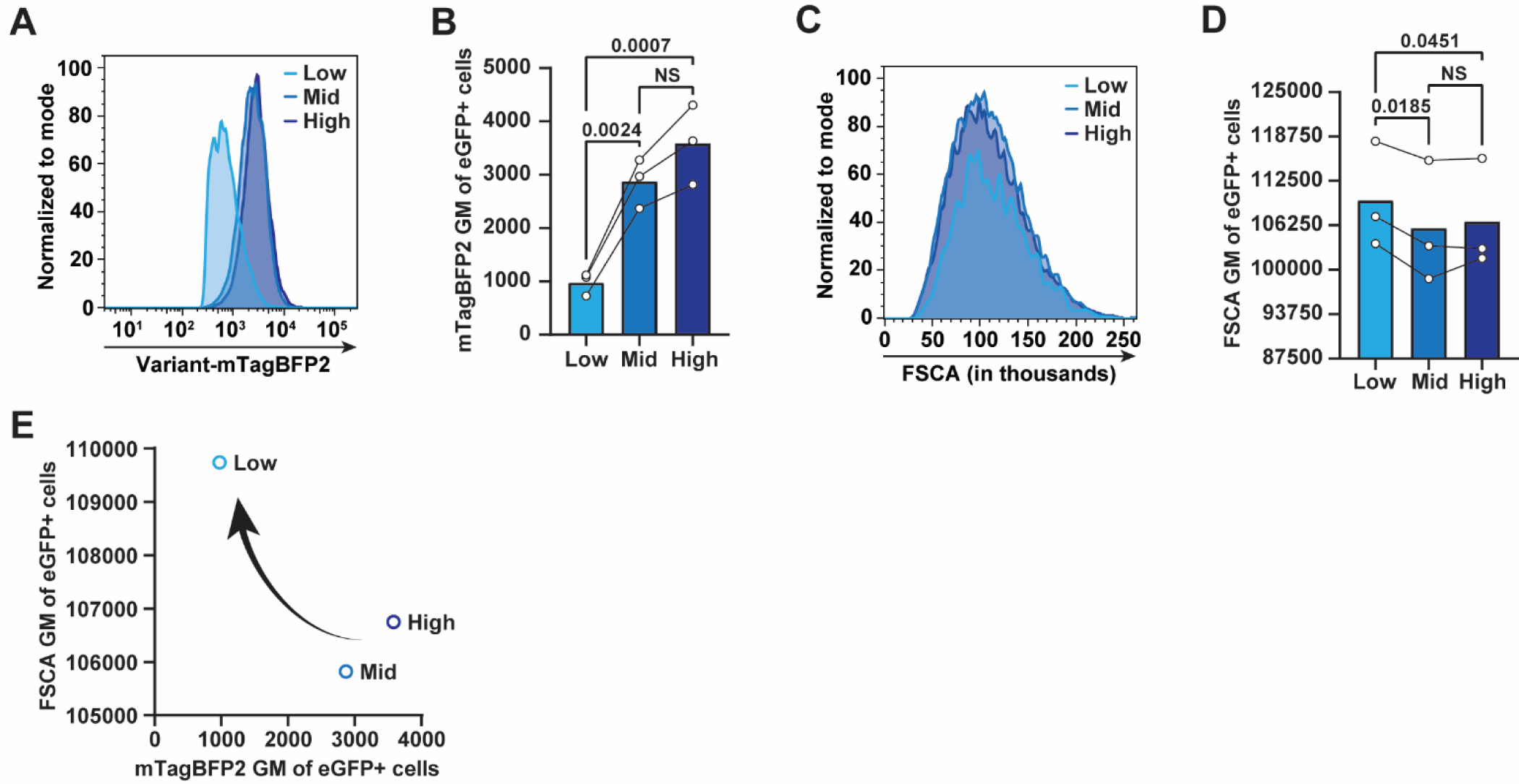
Pooled analyses of *MYH7* variant hiPSC-CM library mTagBFP2 intensity and FSCA, associated with Fig. 2. **A** Representative flow cytometry histogram overlaying mTagBFP2 intensities of MYH7^WT-eGFP/Variant-mTagBFP2^ eGFP^+^ hiPSC-CMs (from Fig. 2) sorted based on low, mid, or high mTagBFP2 intensity and **B** quantifications (*n* = 3 independent replicates). **C** Representative flow cytometry histogram overlaying FSCA values (from Fig. 2) and **D** quantifications (*n* = 3 independent replicates). *P* values calculated using repeated measure one-way ANOVA; *p* < 0.05 considered significant; not significant (NS). **E** Comparison of mean mTagBFP2 GM (x-axis) to mean FSCA GM (y-axis) (from **B** and **D**).

**Supplemental Figure S3.**
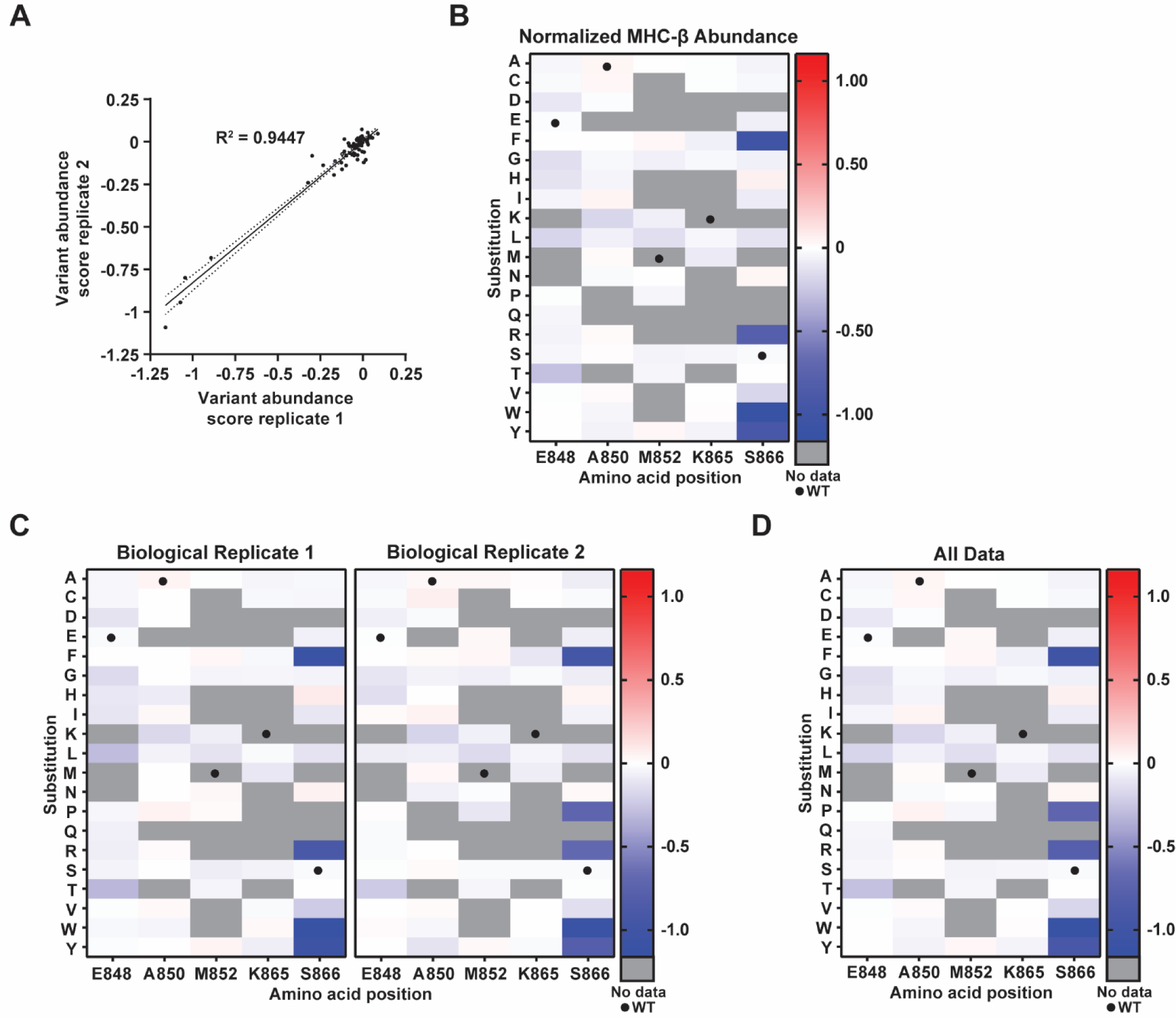
VAMP-seq replicate data, associated with Fig. 2. **A** Simple linear regression of MHC-β-mTagBFP2 abundance scores from replicate 1 (x-axis) relative to replicate 2 (y-axis). Solid line = best-fit line; dotted lines = 95% confidence intervals. **B** Position maps displaying actual MHC-β-mTagBFP2 abundance scores for each mutagenized residue (vertical) relative to amino acid substitutions (horizontal). Grey = no data. Black dots = WT residues. **C** Position map displaying normalized MHC-β-mTagBFP2 abundance scores for biological replicate 1 and 2 (*n* = 1 independent replicate). **D** Position map displaying all MHC-β-mTagBFP2 abundance scores (71 amino acid substitutions) from either replicate (*n* = 1-2 independent replicates). Grey = no data. Black dots = WT residues.

**Supplemental Figure S4.**
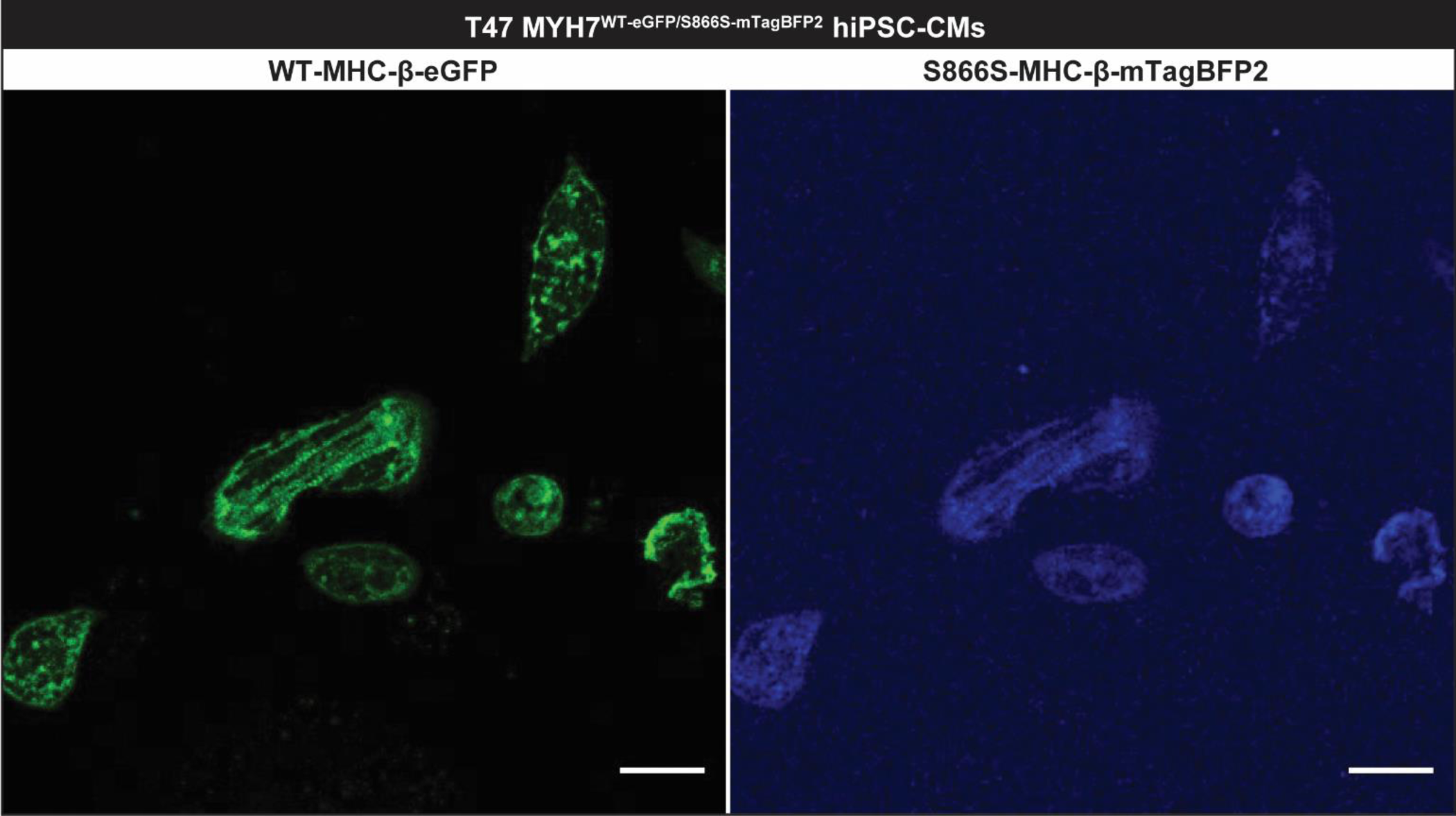
Confocal microscopy of clonal heterozygous *MYH7* variant hiPSC-CMs, related to Fig. 3. **A** Confocal micrograph (60x) of live, T47 cardiomyocytes differentiated from clonal MYH7^WT-eGFP/S866S-mTagBFP2^ hiPSCs. MHC-β-eGFP (green) and MHC-β-mTagBFP2 (blue) expression shown in left and right images, respectively. T = days after the activation of differentiation. Scale bar = 25 µm.

**Supplemental Figure S5.**
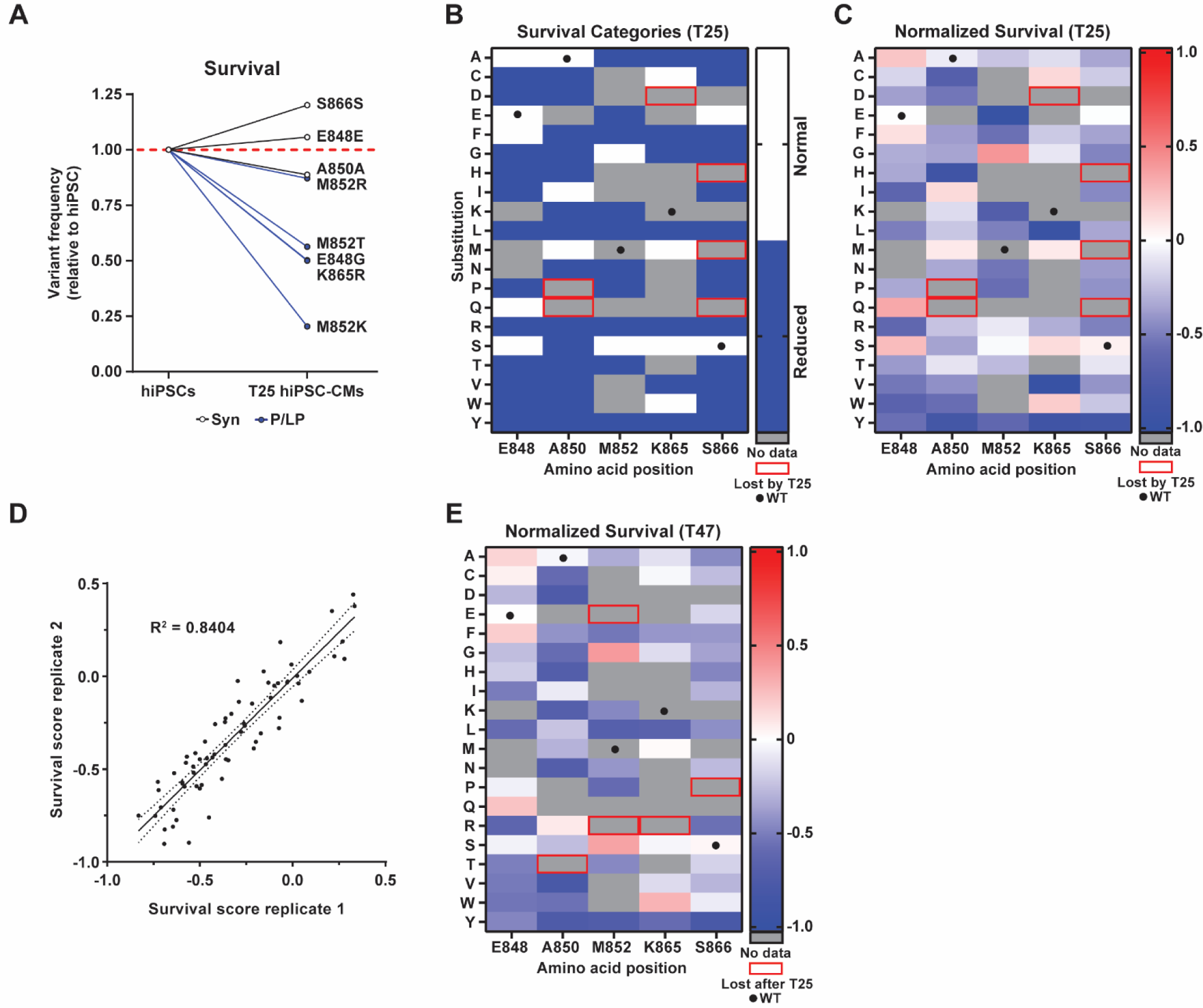
T25 *MYH7* variant hiPSC-CM survival and T47 *MYH7* variant hiPSC-CMs replicate survival data, associated with Fig. 4. Variant frequency of MYH7^WT-eGFP/Variant-mTagBFP2^ hiPSC-CMs with single synonymous (Syn; white) or pathogenic/likely pathogenic (P/LP; blue) *MYH7* variants assessed at T25 (*n* = 1 replicate) relative to starting frequencies as hiPSCs (*n* = 1 replicate) (hiPSC and T25 datasets from Friedman *et al.* [19]). T = days after the activation of differentiation. Position maps displaying T25 normalized categorical (**B**) and actual (**C**) survival scores (relative to hiPSCs) for each mutagenized residue (vertical) relative to amino acid substitutions (horizontal). Grey = no data. Red box = present in hiPSCs but ‘Lost’ at T25. Black dots = WT residues. **D** Simple linear regression of normalized T47 survival scores from replicate 1 (x-axis) relative to replicate 2 (y-axis). Solid line = best-fit line; dotted lines = 95% confidence intervals. **E** Position map displaying actual T47 normalized survival scores (relative to hiPSCs) for each mutagenized residue (vertical) relative to amino acid substitutions (horizontal). Grey = no data. Red box = present in T25 hiPSC-CMs and hiPSCs but ‘Lost’ at T47. Black dots = WT residues. Dashed red line in **A** indicates normalized starting variant frequencies in hiPSCs.

